# Association of seminal polyaromatic hydrocarbons exposome with idiopathic male factor infertility: A proteomic insight into sperm function

**DOI:** 10.1101/2022.03.21.484927

**Authors:** Jasmine Nayak, Soumya Ranjan Jena, Sugandh Kumar, Sujata Kar, Anshuman Dixit, Luna Samanta

## Abstract

Oxidative stress (OS) is implicated in 80% of idiopathic male infertility (IMI) patients where exposure to redox active environmental toxicants such as polyaromatic hydrocarbons (PAHs) may play. In the present study the seminal exposome of various PAH was analyzed in two separate cohorts including 43 fertile donors and 60 IMI patients by HPLC and receiver operator characteristic curve was applied to find out the cut-off limits. Furthermore, spermatozoa from both the groups were subjected to label free liquid chromatography mass spectroscopy (LC-MS/MS) followed by bioinformatics analysis to elucidate the molecular mechanism(s) involved and the key proteins from the affected pathways were validated by western blot along with oxidative modification of proteins. Of the 16 standard PAH 13 were detected in the semen. Receiver Operator Character (ROC) Curve analysis (AUC_ROC_) revealed the PAHs having most significant effect on fertility are of the following order Anthracene<benzo(a)pyrene<benzo[b]fluoranthene<Fluoranthene<benzo(a)anthracene<indol (123CD)pyrene<pyrene<naphthalene<dibenzo(AH)anthracene<fluorene<2bromonaphthalene <chrysene<benzo(GH1)perylene. Benzo[a] pyrene is invariably present in all infertile patients while naphthalene is present in both fertile and IMI group. Of the total 773 detected proteins (Control: 631 and PAH: 717); 71 were differentially expressed (13 underexpressed, 58 overexpressed) in IMI patients resulting in impaired mitochondrial dysfunction and oxidative phosphorylation, DNA damage, Aryl hydrocarbon receptor (AHR) signalling, xenobiotic metabolism and induction of NRF-2 mediated oxidative stress response (increased in 4-hydroxynonenal and nitrosylated protein adduct formation, and declined antioxidant defence). The increased GSH/GSSG ratio in patients may be an adaptive response to metabolize the xenobiotics via conjugation as evidenced by overexpression of AHR and Heat shock protein 90 beta (HSP90β) in patients. Seminal PAH concentrations, oxidative protein modification along with protein markers (e.g. AHR and HSP90β) may help in better prediction and management of IMI. Contribution of environment borne PAH in semen should not undermined in infertility evaluation.

## 1. Introduction

Semen quality of men in their reproductiveage is markedly deteriorating over past decades^1,2^. Approximately 15% of the co-inhabiting couples are infertile where 50% of them have abnormal semen parameters implying the involvement of male-infertility-associated factor ^3^.Male infertility in general is known as a multi-causal effect with a very few large-scale epidemiological reports available^4,5^.Due to the paucity of information on causative factors ondecline in semen quality; accurate diagnosis and personalized treatment options are restricted. An infertility case withunknown causative factor is identified and referred to as idiopathic infertility^3,6^. It is reported that ∼75% of oligospermic men are idiopathic^7^. Albeit, the interplay between genetic, environmental and lifestyle factors are proposed to be behind this condition; very few reports established the role of environmental toxins in idiopathicinfertility^8^.Most of the studies simply establish a correlation between environmental toxin levels in body fluids such as blood plasma or urine with semen parameter without giving much insight into the mechanisms involved ^8,9^.European Association of Urology attributed idiopathic male factor infertility to endocrine disruption due to environmental pollution, reactive oxygen species, or genetic abnormalities ^10^. In recent times, more emphasis is given to the “exposomes” concept that refers to the totality of environmental exposures of an individual during the lifetime. This novel approach combines the body burden of environmental toxins and modern omics technologies to study the role of the environment in human diseases^11^.Many environmental toxinssuch as pesticides, herbicides, phthalates and polyaromatic hydrocarbons (PAHs) undergo metabolic activation in human body and cause oxidative stress^12^. Oxidative stress is time and again reported to not just correlate with defective sperm function but is causally involved in the genesis male factor infertility^13-15^. Male Oxidative Stress Infertility (MOSI) is proposed for the management of idiopathic male infertility with measurement of seminal oxidation-reduction potential (ORP) as an easy clinical biomarker^16^. It is further suggested that upto 80% of the total cases of idiopathic infertility have augmented oxidative stress^13-15^.Therefore, it is imperative to look into the cause behind the aetiology of MOSI in idiopathic infertility as a function of environmental toxins.

The crucial environmental toxins, especially phthalates, bisphenols, pesticides, flame retardants and Polyaromatic hydrocarbons (PAHs) warrant special attention due to their potential role as endocrine disruptors affecting hypothalamo-pituitary-thyroid axis and hypothalamo-pituitary-gonadal axis^17^. PAHs are the by-products of incomplete combustion of organic materials generated from tobacco and cigarette smoke, barbequed food, vehicle exhaust and oil spillers as well as during coke production and chemical manufacturing. They are usually metabolically activated by cytochrome *P450* enzymes during steroidogenesis and promotes free radical generation ^18^. The ROS (H_2_O_2_and O_2_ ^·−^) generated during normal steroidogenesis are within critical levels and play an important role in the regulation of steroidogenic activity of the Leydig cell^19^. The elevated production of ROS have been found to inhibit steroid productions, and causes damage to mitochondrial membrane of spermatozoa^20^.However, our knowledge on oxidative stress-induced idiopathic male infertility as a function of environmental borne seminal concentration of PAH is extremely limited in general and with respect to non-occupational exposure in particular. With this background, the present study in designed to analyse the level of PAH in the ejaculate of idiopathic infertile men and its relationship with induced oxidative stress via high throughput shotgun proteomic analysis in comparison to proven fertile donors to unravel the pathways involved towards discovery of plausible biomarkers.

## 2. Materials & Methods

### 2.1 Ethics statement and Patient selection

Patients attending the infertility centre and the proven fertile donors at Kar Clinic and Hospital Pvt. Ltd., Bhubaneswar, Odisha, India were recruited for the studyafter approval by the Institutional Ethics Committee. All participants gave an informed written consent to be included in this study.The exclusion criteria were leukocytospermia (Endtz positive), azoospermia, historyof systemic illness, inflammation of reproductive tract (orchitis, epididymitis, urethritis, and testicular atrophy), sexually transmitted disease, and medications.The participants included in this study were non-smokers, non-alcoholic, and had a normal body mass index. Healthy donors (with no known medical condition) who had established fertility recently i.e., within one year with no cases of embryo loss were included as the control group.Subsequently upon estimation of PAH concentrations and receiver operator characteristic (ROC) curve analysis (as described below) in comparison to fertile donors, patients were segregated into PAH positive infertile group.Of the total 60 patients recruited for the study, 18 idiopathic infertile patientswere excluded based on their lower than cut off value of one or more PAH present in their semen. Therefore, in the final step 43 proven fertile donor were compared with 42 PAH positive idiopathic infertile patients.

### 2.2 Semen analysis

The semen samples were collected from all participants of both groups (idiopathic infertile patientsn=60 and fertile donor n=43) by masturbation after 3-5 days of sexual abstinence. Samples were allowed to liquefy completely for 20-30 min at 37°C followed by semen analysis as per World Health Organization (WHO) 2010 guidelines (WHO 2010). Basic semen analysis included both macroscopic (volume, pH, colour, viscosity, liquefaction time) and microscopic parameters such as sperm concentration, motility and morphology, as well as peroxidase or Endtz test. Samples with >1.0 × 10^6^/ml round cell with a positive peroxidase test were excluded from the study.After liquefaction and semen analysis, the samples were subjected to centrifugation at 400 g for 20 min at 37°C to separate the sperm and seminal plasma. The seminal plasma was processed for PAH measurement.

### 2.3 Measurement of seminal PAH exposomes

The HPLC analysis was carried out by injecting 20 μL of the seminal plasma into the chromatographic system (Thermo Scientific UltiMate 3000) for the determination of PAHs concentration using PAH standards. PAHs were segregated by C-18 column with a gradient elution process using solvent water and acetonitrile. The elution conditions applied were: 0 – 20 min, 40% of acetonitrile isocratic; 20 – 37 min, 50-100% of acetonitrile gradient, 37 – 42 min, 100% of acetonitrile isocratic, 42 – 45 min, 100 - 40% of acetonitrile, gradient. The flow rate was set at 1 mL/min, at room temperature. Under these conditions, PAHs could be separated satisfactorily within 45 min. The PAHs were identified by comparing the retention time with those of standards taken. The concentration of PAH in semen was calculated according to the formula:

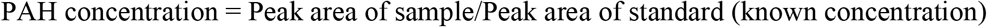

### 2.4 Determination of PAH threshold for incidence of infertility

Receiver operating characteristics (ROC) curve is used in clinical biochemistry for the determination of the cut-off point in clinical deterioration. A ROC curve shows a graphical plot illustrating the diagnostic ability of a binary classifiers to identify the sensitivity and the specificity of PAH levels in the prediction of male fertility status. The ROC curves were createdusing total individual PAHs as ‘‘test variables,’’ and ‘‘fertile donor” versus idiopathic infertile patients’’ (binary variable, with fertile ¼ 0 and infertile patient ¼ 1) as ‘‘state variable’’ and setting the value of the ‘‘state variable’’ as 1. The optimal cut-off value was determined with the use of the Youden index to maximize the sum of sensitivity and specificity.

### 2.5 Sperm protein Extraction and estimation

Post separation from seminal plasma, sperm pellet extracted was washed thrice with phosphate buffer saline (PBS) and centrifuged at 400 g for 10 min, at 4°C. Sperm lysate was prepared by adding 100μl of Radio-immunoprecipitation assay (RIPA) buffer supplemented with Protease inhibitor cocktail (cOmplete ULTRA Tablets; Roche) to the sperm pellet and left overnight at 4 °C for complete cell lysis. The lysate was centrifuged at 14,000 g for 30 min at 4 °C followed by separation of the supernatant. Protein quantification of the supernatant was determined using bicinchoninic acid (BCA; Thermo Fisher Scientific, Waltham, MA).

### 2.6 Assessment of Glutathione and Redox potential

The total GSH equivalents (GSH + GSSG) were spectrophotometrically quantified by glutathione reductase (GR) recycling assay at the expense of oxidation of NADPH using 5-5’-dithiobis 2-nitrobenzoic acid (DTNB; Ellman’s reagent). Similarly, GSSG was measured by masking GSH with 2-vinylpyridine. Formeasurement of the total GSH equivalents (GSH+GSSG), GR was added to the assay mixture for reduction of GSSG to GSH at the expense of oxidation of NADPH. The reduction potential (Ehc) of GSH/GSSG couple was calculated by using Nernst equation. The sperm lysate (described above) was precipitated with ice-cold 5% trichloroacetic acid containing 0.01N HCl, and cleared by centrifugation. The deproteinized supernatants were used for the assay. In brief, the assay mixture (final volume 200 μl) contained 3 mM NADPH in 125 mM Phosphate buffer containing 6.3 mM EDTA (pH 7.5), DTNB (0.6 mM) and sperm lysate (25 μg protein). To this 2 μl GR (∼1 Unit, Sigma-Aldrich, St. Louis, MO, USA) was added and the yellow chromatophore (2-nitro-5-thiobenzoate: TNB2-) formed by the interaction of SH groups from GSH and GSSG (after conversion by GR) with DTNB was recorded at 405 nm in a iMark Absorbance Microplate Reader (BioRad Instruments, Inc., Japan) at 1-min intervals for 6 min. All the determinations were normalized to protein content. The absolute GSH amount was quantified from difference between the total GSH equivalent and the obtained GSSG value.

The glutathione redox potential (E^hc^) was calculated by Nernst equation for half reaction:

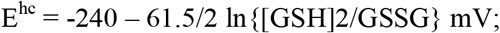

where −240mV is the standard redox potential (E°) of GSH at pH 0, −61.5/2 denotes RT/z F i.e R= Gas constant (8.314 J K-1 mol-1), T= absolute temperature of 37°C or 310 K, F= Faraday constant (9.64853 × 104 C mol-1), z = number of electrons exchanged in the chemical reaction GSSG + 2e-+ 2H → 2GSH.

### 2.7 Shotgun Proteome profiling of spermatozoa

The extracted spermatozoa proteins were subjected to LC-MS/MS for proteomic analysis. 50 μl of sperm lysate from each sample (n=3 per group; one pooled sample and two individual samples) were taken and reduced with 5 mM TCEP, then alkylated with 50 mM iodoacetamide and subsequently digested with Trypsin (1:50, Trypsin/lysate ratio) for 16 h at 37 °C. To eliminate the salt, these digests were cleansed using a C18 silica cartridge and then dried with a speed vac. Then the dried pellet was resuspended in buffer A (5% acetonitrile, 0.1% formic acid). All the experiments were carried out using an EASY-nLC 1000 system (Thermo Fisher Scientific) combined with Thermo Fisher-QEXACTIVE mass spectrometer designed with nano-electrospray ion source. A 25 cm PicoFrit column (360μm outer diameter, 75μm inner diameter, 10μm tip) packed with 1.8 μm of C18-resin (Dr Maeisch, Germany) was employed to resolve 1.0 μg of the peptide mixture. The peptides were loaded with buffer A and eluted at a flow rate of 300 nl/min for 100 minutes with a 0–40% gradient of buffer B (95% acetonitrile, 0.1% formic acid). The acquisition of MS data is done by considering the most abundant precursor ions from the survey scan using top 10 data-dependent method.

### 2.8 Data Processing

All the samples were processed and RAW files generated were analyzed with comparison to Uniprot HUMAN reference proteome (HUPO) database using Proteome Discoverer (v2.2). The precursor was set at 10 ppm and fragment mass tolerances was set at 0.5 Da for SEQUEST search. The protease used to produce peptides, where enzyme specificity was set for trypsin/P (cleavage at the C terminus of “K/R: unless followed by “P”) along with maximum missed cleavages value of two. For database search, carbamidomethyl on cysteine was considered as fixed modification while N-terminal acetylation and oxidation of methionine were considered as variable modifications. Both protein False Discovery Rate (FDR) and peptide spectrum match were set to 0.01.

### 2.9 Quantitative proteomics

The following procedures were performed to execute Label free quantification (LFQ) suit for quantitative proteomics using MaxQuant v1.5.2.8 (http://www.maxquant.org/): feature identification, initial Andromeda search, recalibration, main Andromeda search, and posterior error probability calculation (likelihood of a protein being incorrectly recognized). At first, razor peptides and protein groups were identified. Proteins that cannot be identified unambiguously by distinct peptides but share peptides were grouped together and quantified as a single protein group. For instance, if all detected peptides of protein X were also identified for protein Y, X and Y are classified as one protein group (even though unique peptides were found for Y, since it was still uncertain whether X was present in the sample). Only one common quantification was generated for both proteins in the result. A Razor peptide is formed when two protein groups (protein A and protein B) are unequivocally identified by distinct peptides yet share a common peptide. Following the discovery of distinct and unique peptides, a “match across the runs” procedure was used to match the same accurate masses across several LC-MS/MS runs within a 1.5-minute retention time frame. Relative quantification was determined by comparing the abundance of the same peptide species/protein across runs, whereas absolute quantification was determined by equating the quantities of various proteins in the same sample. The MaxQuant algorithms use peak detection and scoring of peptides, as well as mass calibration and protein quantification, to provide summary statistics. Protein abundances were normalized using the LFQ algorithm in MaxQuant and then converted to Log_2_ for further analysis. The label-free method analyses the intensity of these peptides to determine peptide ratios by taking the greatest number of detected peptides between any two samples. The median values of all peptide ratios of a given protein are used to calculate protein abundance.

### 2.10 Bioinformatics analysis

The differentially expressed proteins (DEPs) calculated based on Log_2_ fold change > 1 and p-value < 0.05 were subjected to functional annotation and enrichment analysis by means of publicly available bioinformatics annotation tools and databases such as String, UniProt, and Cytoscape. Enriched terms were ranked by p-value (hypergeometric test) using Cytoscape ClueGO plugin. Venn diagram showing distribution pattern of proteins were drawn using Venny 2.1. Hierarchical clustering of the DEPs between fertile donor and idiopathic infertile patients were analysed by the construction of heat map using R.3.4.4 package (Complex Heatmap map library). Euclidean distance correlation matrix was used for hierarchical clustering of the DEPs for dendrogram plotting showing complete linkage between the proteins. The Biological Networks Gene Ontology (BiNGO) application in Cytoscape was used for the determination of significantly overrepresented Gene Ontology (GO) terms in the DEPs data set and the predominant functional themes of the tested DEPs were mapped to visualize the biological pathways altered in the infertile group. The protein-protein networks were obtained from the STRING database (http://string-db.org/). A p < 0.05 was considered significant. Proprietary curated database such as Ingenuity pathway analysis (IPA) was used to analyze the involvement of DEPs in biological and cellular processes, pathways, cellular distribution, protein-protein interactions and regulatory networks.

### 2.11 Western blotting

Two key protein markers of PAH metabolism, i.e., AhR and HSP90β(sc-101104,sc-59578, Santacruz, Mouse)were validated by western blotting. Besides, the impact of induced oxidative stress on protein modifications, namely, 4-Hydroxynonenal (HNE)(ab46545, Rabbit, Abcam) protein adduct formation and anti-3-nitro-tyrosine(ab110282, Mouse, Abcam) were also studied by western blotting. From every group two individual and one pooled samples were run in duplicates to maintain biological and technical variabilities. Samples were normalized for protein concentration in each group. Washed spermatozoa were lysed in RIPA lysis buffer (Sigma-Aldrich, St. Louis, MO, USA) overnight at 4ºC containing proteinase inhibitor cocktail (Roche, Indianapolis, IN, USA). Samples containing 20-30 μg of protein in 15 μl volume per sample were separated on a 4-20% SDS-PAGE and electroblotted onto polyvinylidenedifluoride (PVDF) membranes. Then the proteins transferred to PVDF membrane was blocked with 5% non-fat dry milk in Tris-buffered saline Tween 20 (TBST) buffer for 2 hrs. After that primary antibody incubation was done overnight (4°C) followed by the specific secondary antibodies (Mouse, Rabbit, Abcam) at room temperature for 3 hrs. Blots were then washed using TBST and protein bands were visualized using an enhanced chemiluminescence kit-Pierce™ ECL Western Blotting Substrate (Thermo Scientific, Rockford, IL, USA) in ChemiDoc™ MP Imaging System (BioRad, Hercules, CA, USA).

The densitometric analysis of western blot images was done through Image J software (NIH, Bethesda, MD) by total intensity normalisation method. Results were expressed as fold change relative to the fertile donor.

### 2.12 Statistical Analysis

Statistical analysis was performed by MedCalc Statistical Software, ver. 17.4 (MedCalc Software; Ostend, Belgium). The data are expressed as mean□±□standard deviation (SD). Normalization of data was assessed using Shapiro–Wilk test followed by Levene’s test for homogeneity of variance. Data on semen parameters and biochemical estimation were analyzed by Mann-Whitney U-test. Results of LC-MS/MS proteomics and Western blotting was subjected to Welch’s t-test, or unequal variances t-test. A difference of p<0.05 (minimum) was considered significant.

## 3. Results

### 3.1 Semen analysis

Seminogram results are presented in **Table S1**. All idiopathic infertile patients included in this study have at least one semen parameter in semen analysis below WHO 2010 criteria. However, the average values were within the range, but the sperm concentration, motility, morphology and vitality were considered significant.

### 3.2 PAH Exposome in semen

Out of the 16 standard PAHs used for screening, a total of 13 PAH metabolites, i.e., Anthracene, Benzo (A) Anthracene, Benzo (A) Pyrene, Benzo (B) Fluoranthene, Benzo (GH1) Perylene, Chrysene, Dibenzo (AH) Anthracene, Fluorene, Fluoranthene, Indo (123 CD) Pyrene, Napthalene, 2 Bromonapthalene, Phenanthrene, Pyrene were detected in the semen samples at ng/ml level **(Fig 1)**. However, the concentration of PAH in the semen of idiopathic infertile patients were significantly higher in comparison to the fertile donor.The cut-off level of these PAHs (if any) determined by ROC curve analysis segregates the fertile from the idiopathic infertile patients. The results of the ROC analysis are presented in **Fig. 1**,**Table S2**. Benzo (A) Pyrene particularly is found to be highly discriminative among 13 PAHs in idiopathic infertile patients.

**Fig. 1.**
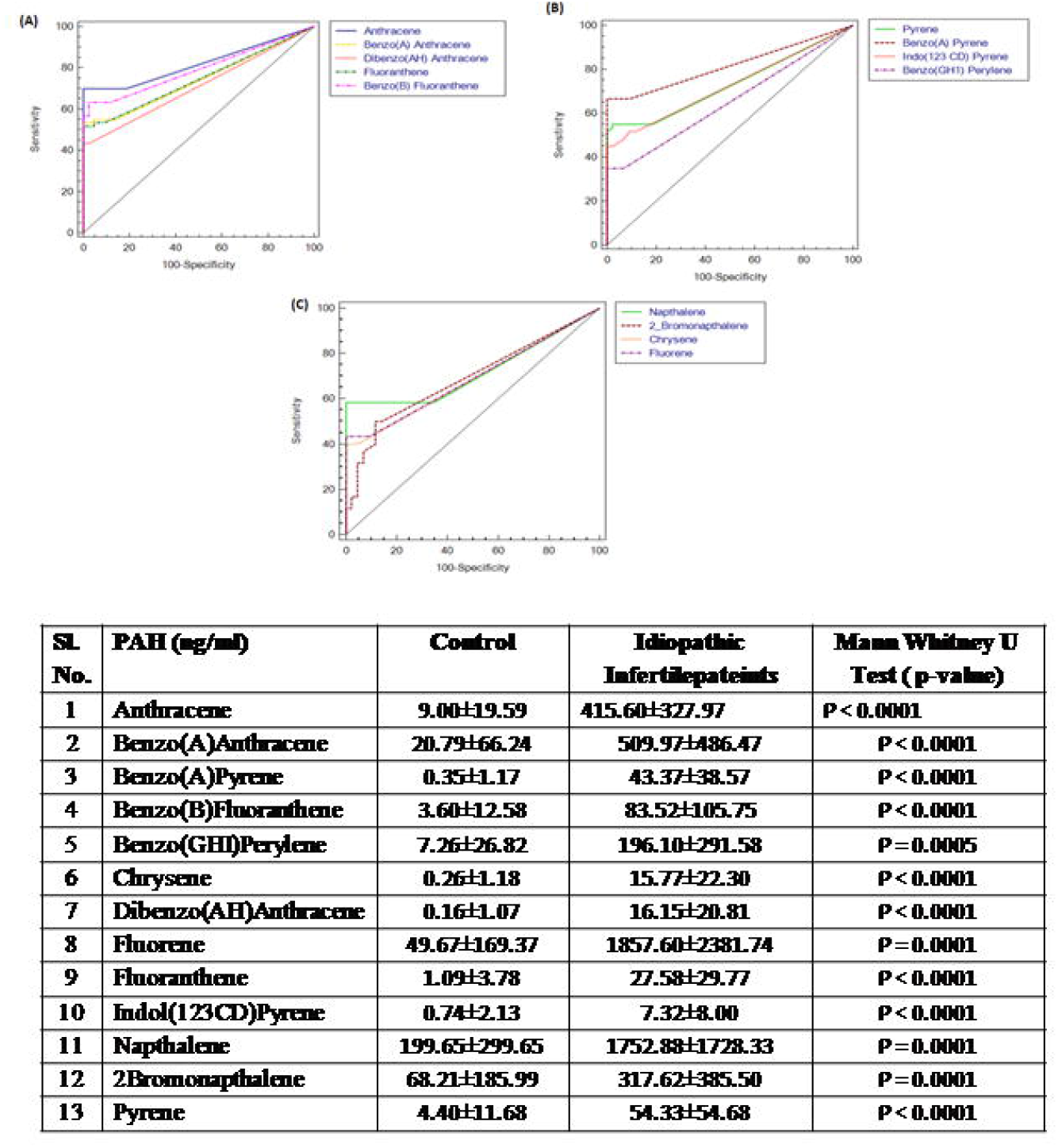
Receiver Operating Characteristic (ROC) curves of polyaromatic hydrocarbons in semen of idiopathic infertile men (n=60) in comparison to fertile donor (n=43).

### 3.3 Effect of PAH concentration on Sperm redox status

To corroborate the alteration in the redox environment of sperm in the idiopathic infertile patients, ratio of GSH:GSSG was measured. An increase in the absolute concentrations of GSH and GSH:GSSG ratio was noticed in idiopathic infertile patients (**Fig. 2A,B**). The reduction potential in spermatozoa of idiopathic infertile patients is more negative with respect to fertile donor (**Fig. 2C**). A reduction in the level of total antioxidant capacity of sperm was observed in the infertile group as compared to fertile control (**Fig. 2D**).

**Fig. 2.**
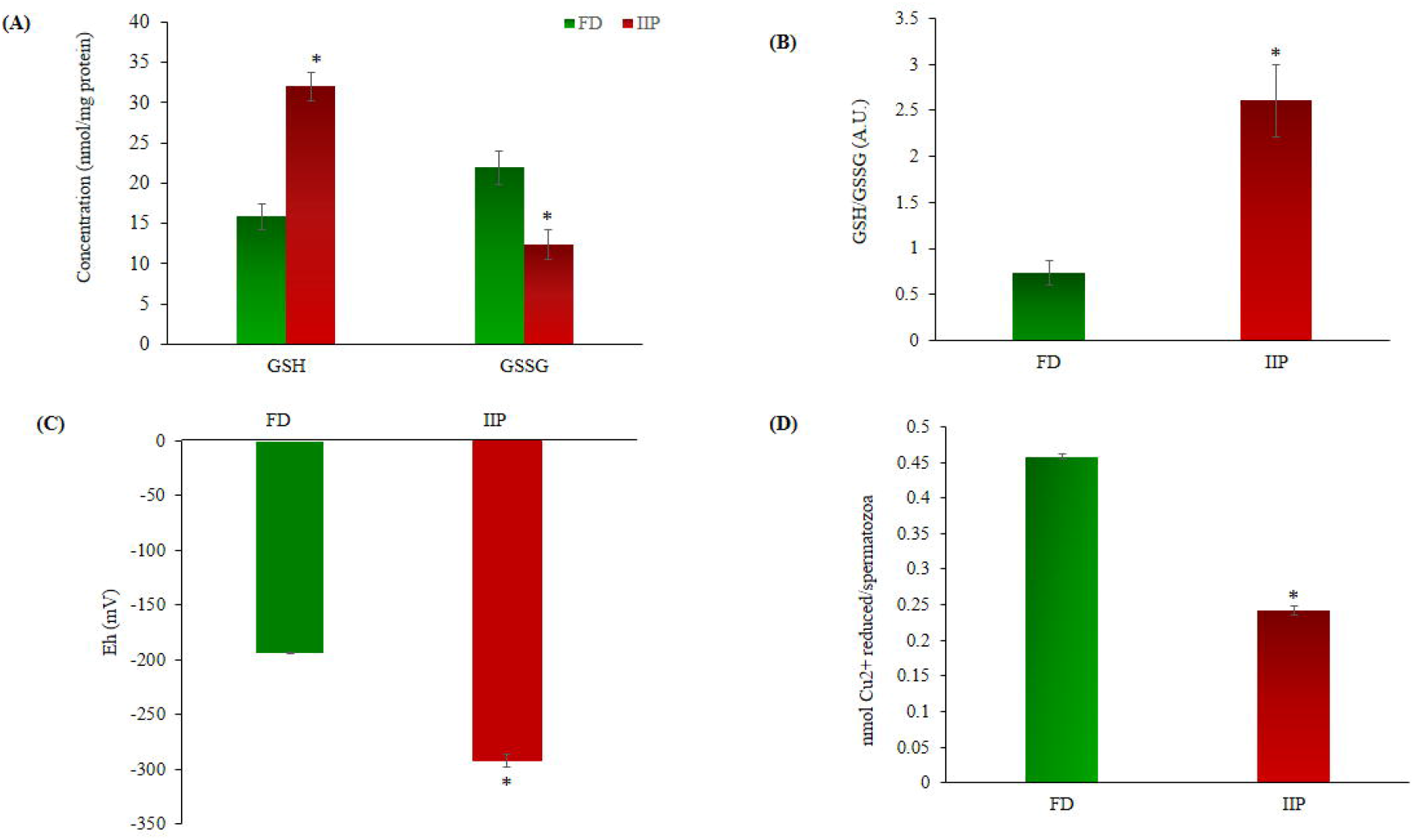
Comparison of redox status in the spermatozoa of fertile donor (n=43) and infertile patients (n=60). A. Levels of reduced (GSH) and Oxidized glutathione (GSSG); B. Spermatozoal redox status; C. Half cell reduction potential. D. Total antioxidant capacity. FD: Fertile donors; IIP: Idiopathic infertility patients. Data are expressed as mean ± SD. *p<0.05.

### 3.4 Global proteome profiling of spermatozoa

The quantitative differential proteomic analysis identified a total of 773 proteins in fertile donor and idiopathic infertile patients by label free LC-MS/MS. Out of the total 773 proteins, 631 were fromfertile donor and 717 from idiopathic infertile patients with 575 proteins common in both **(Fig. 3)**. A total of 71 DEPs (based on Log_2_ fold change >1 and p-value < 0.05) were detected, of which 13 and 58 were under- and over-expressed, respectively in idiopathic infertile patients compared to fertile donor **(Fig. 3, Table S3)**. The hierarchical clustering by heat map showed over and under expressed proteins into the two groups distinctively **(Fig. 3)**.

**Fig. 3.**
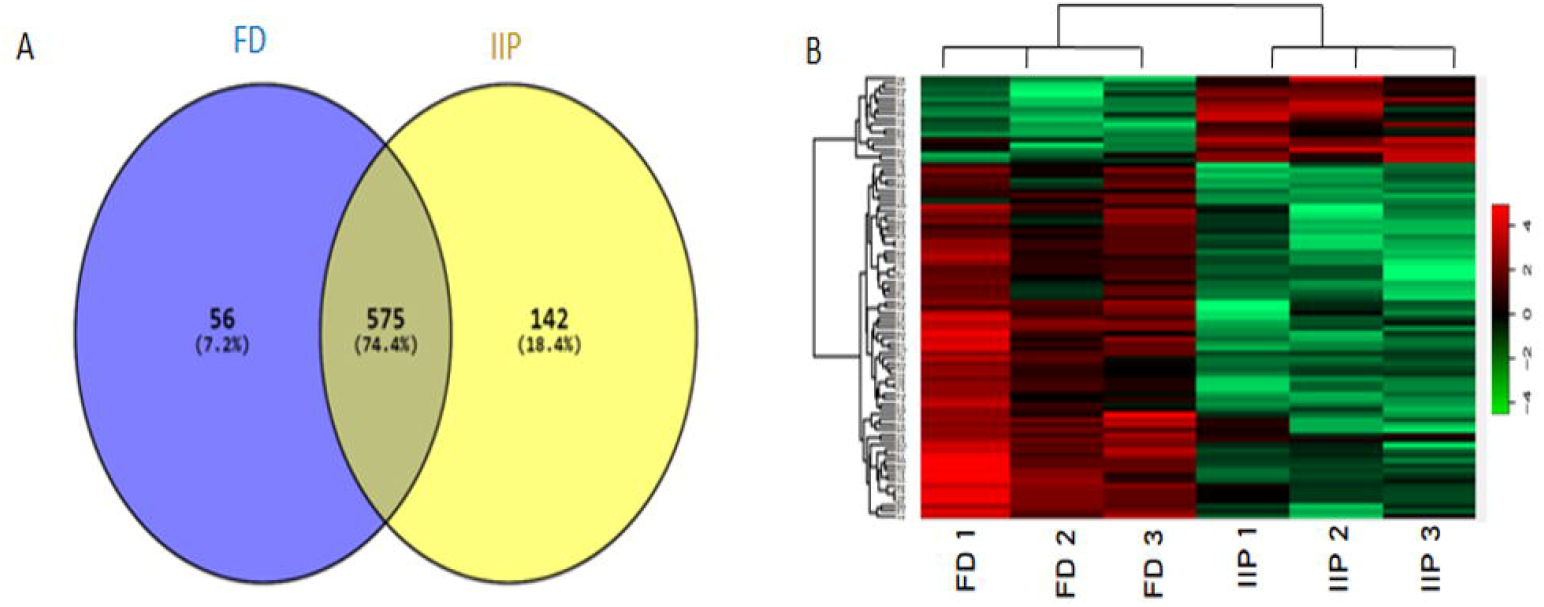
Comparative global proteomic profiling of spermatozoa between FD: Fertile Donors and IIP: Idiopathic infertility patients. (A) Venn diagram showing distribution of differentially expressed proteins (DEPs) (B) Heat map showing a hierarchical cluster of DEPs. The dendrogram for sample replicates (column clustering) separated the samples according to their clinical diagnosis into FD and IIF. Hierarchical clustering analysis between protein expression profiles of DEPs (row clustering) separated overexpressed DEPs in IIP from underexpressed DEPs in FD. The green and red colour denoted low and high expression levels respectively as shown in attached graduated colour scale bar.

The functional enrichment analysis of Gene Ontology (GO) by ClueGO revealed that the identified proteins were involved in various crucial biological functions such as chromosome condensation (GO:0030261), hexokinase activity (GO:0004396), nucleosome assembly (GO:0006334), canonical glycolysis (GO:0061621) and NADH regeneration (GO:0006735).The enriched cellular component and molecular functions were ATPase dependent transmembrane transport complex (GO:0098533), sperm flagellum (GO:0036126), endocytic vesicle lumen (GO:0071682), nucleosome (GO:0000786), metallo-exopeptidase activity GO:0008235, hexokinase activity (GO:0004396) and glucose binding (GO:0005536).The enriched processes and the identified proteins involved in various molecular functions along with its localisation were shown in **Fig. 4, Table S4**.

**Fig. 4.**
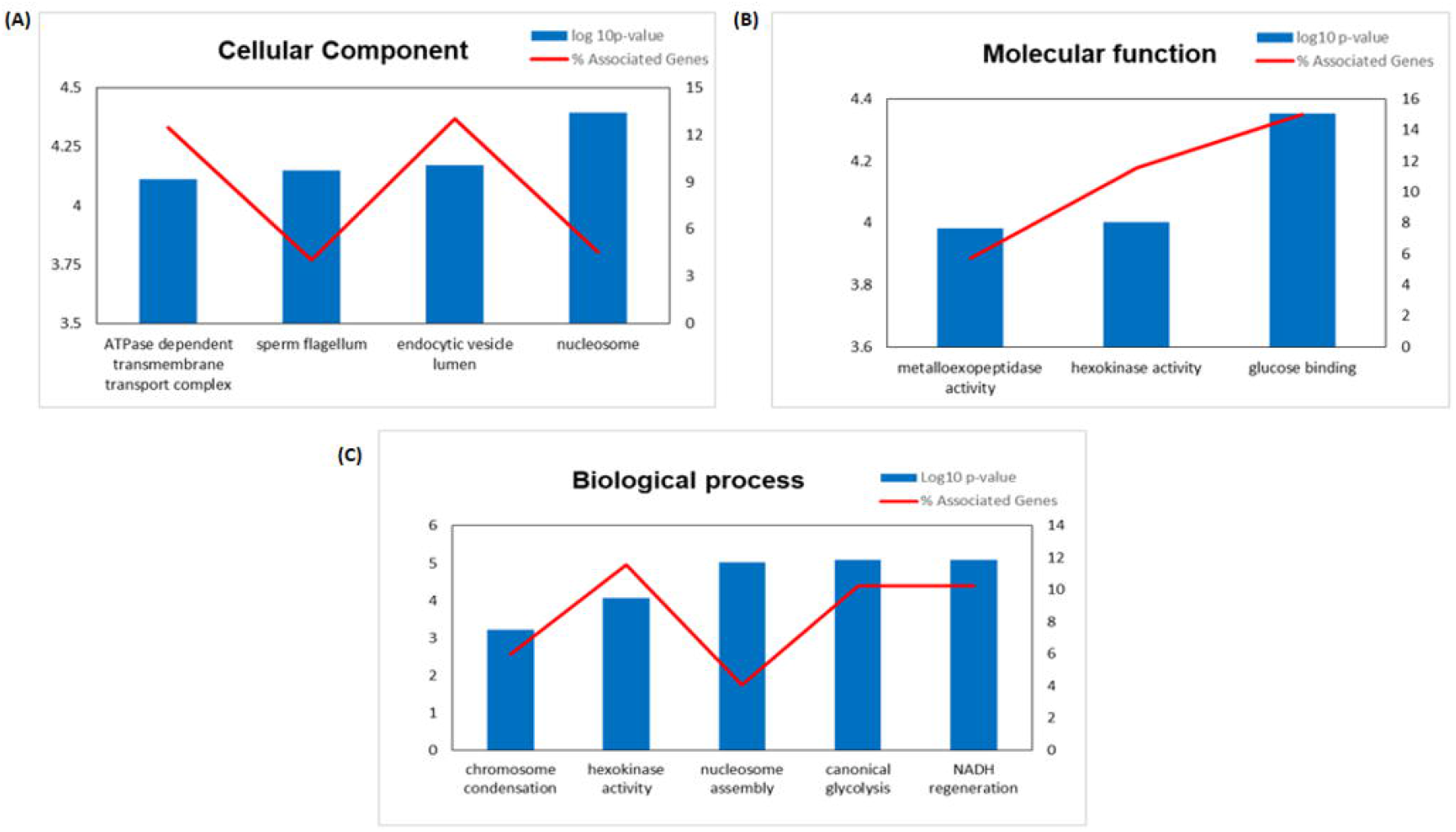
Gene Ontology (GO) enrichment analysis result of differentially expressed proteins (DEPs) in IIP: Idiopathic infertile patients compared to FD: fertile donor. Bar graph showing the top GO terms for cellular component (A) molecular function (B) and biological process (C).

### 3.5 Functional pathway analysis of Differentially Expressed Protein

The BiNGO mapping revealed the involvement of DEPs in reproduction, spermatogenesis, nucleosome assembly, chromatin assembly, DNA packaging and glycolysis (Bonferroni step down with p value ≤0.001) resulting in DNA damage, impaired energy metabolism and reproductive function **(Fig. 5)**. String protein-protein interaction analysis of DEPs revealed that the major pathways deregulated are Glycolysis/Gluconeogenesis (HAS:00010; FDR 1.15e-06), Fatty acid degradation (HAS:00071; FDR 0.0054),HIF-1 signaling pathway (HAS:04066; FDR 0.0281), Estrogen signaling pathway (HAS:04915; FDR 0.0498), Oxidative phosphorylation (HAS:00190; FDR 0.0498), Metabolic pathways (HAS:01100; FDR 0.0029),DNA packaging (GO:0006323; FDR 0.00072),Regulation of regulation of reactive oxygen species metabolic process (GO:2000377; FDR 0.0414), Post-translational protein modification (GO:0043687;FDR 0.0080), and Spermatogenesis (GO:0007283; FDR 0.0284) **(Fig. S1)**. The protein interaction of upregulated DEPs by IPA identified the topmost molecular network to be associated with Cancer, Endocrine System Disorders, Organismal Injury and Abnormalities where out of the 35 nodal proteins 18 were detected in our dataset. In the second most pathway the proteins were involved in Cell Death and Survival, Cellular Development, Organismal Survival where out of 21 nodal proteins 12 were from our dataset. The downregulated DEPs topmost network is associated with Cancer, Cell Death and Survival, Organismal Injury and Abnormalities where out of 19 nodal proteins 9 were found in our dataset **(Fig. 6 & 7, Table S5)**.

**Fig. 5.**
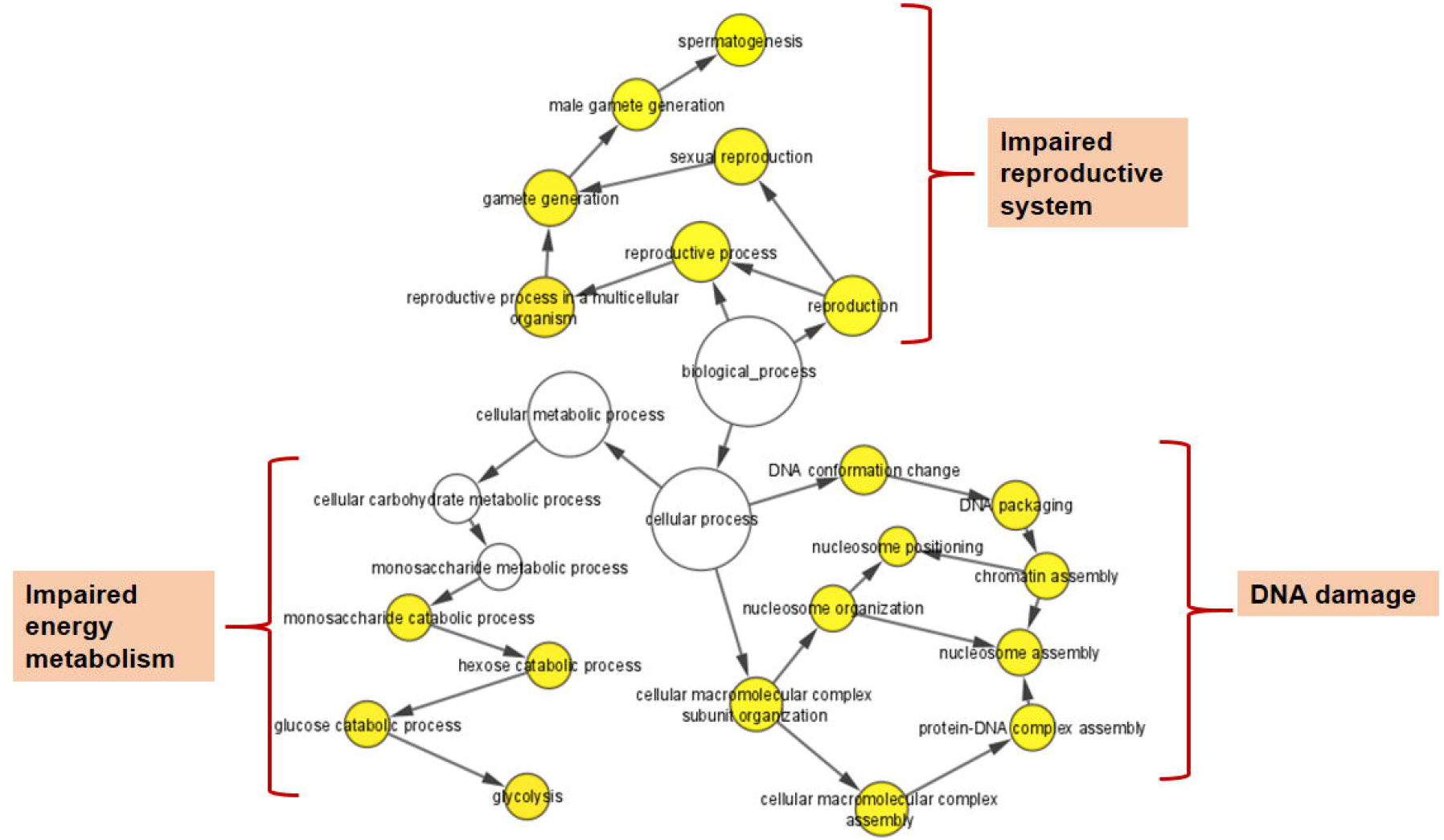
Cytoscape (BiNGO app) enrichment analysis revealed over-represented biological processes for the differentially expressed proteins (DEPs) in the spermatozoa of idiopathic infertile patients in comparison to fertile donor.

**Fig. 6.**
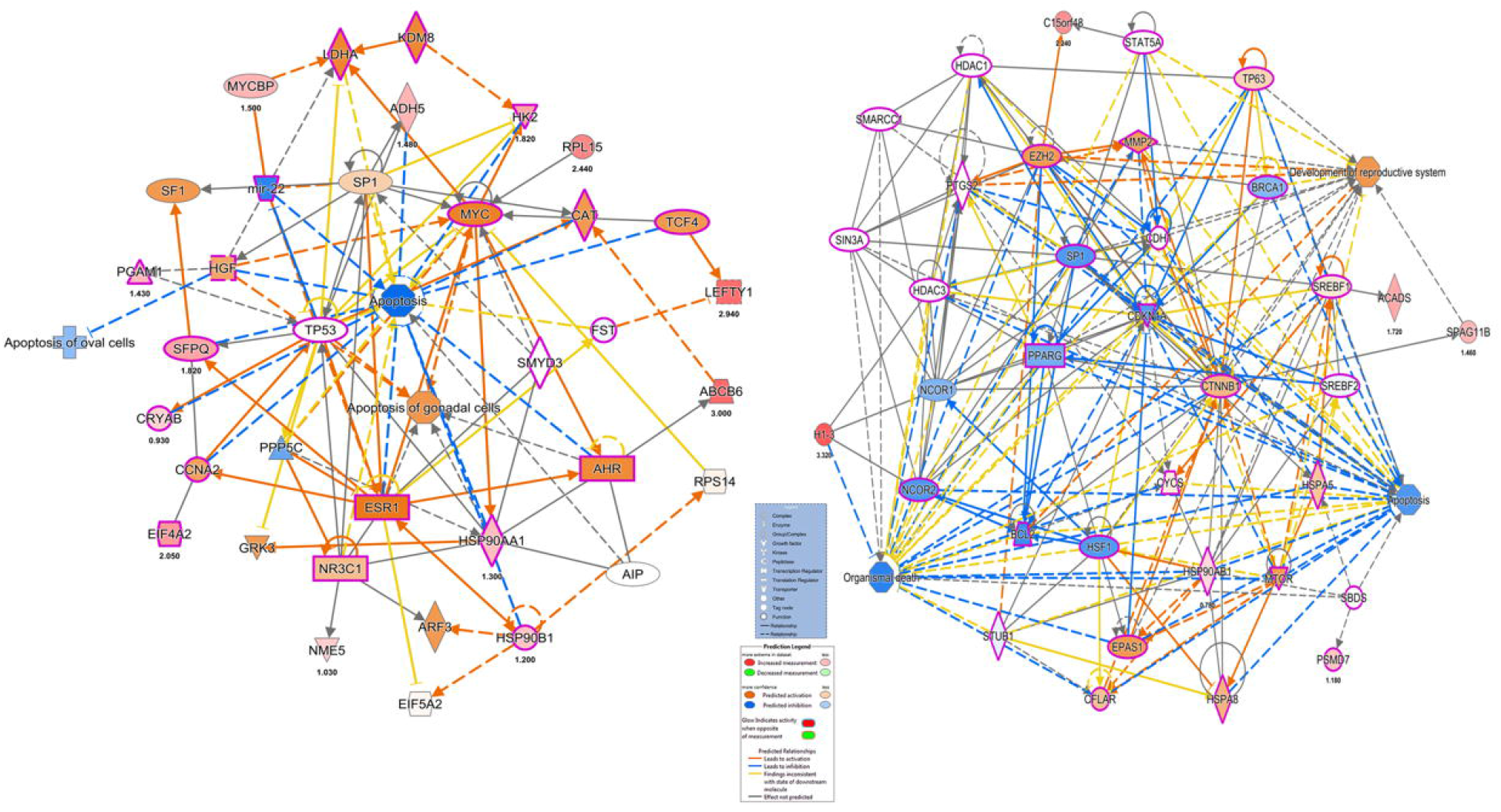
Ingenuity pathway Analysis of overexpressed proteins in Idiopathic infertile patients compared to fertile donor top Disease and Function (A) Cancer, Endocrine System Disorders, Organismal Injury and Abnormalities (B) Cell Death and Survival, Cellular Development, Organismal Survival.

**Fig. 7.**
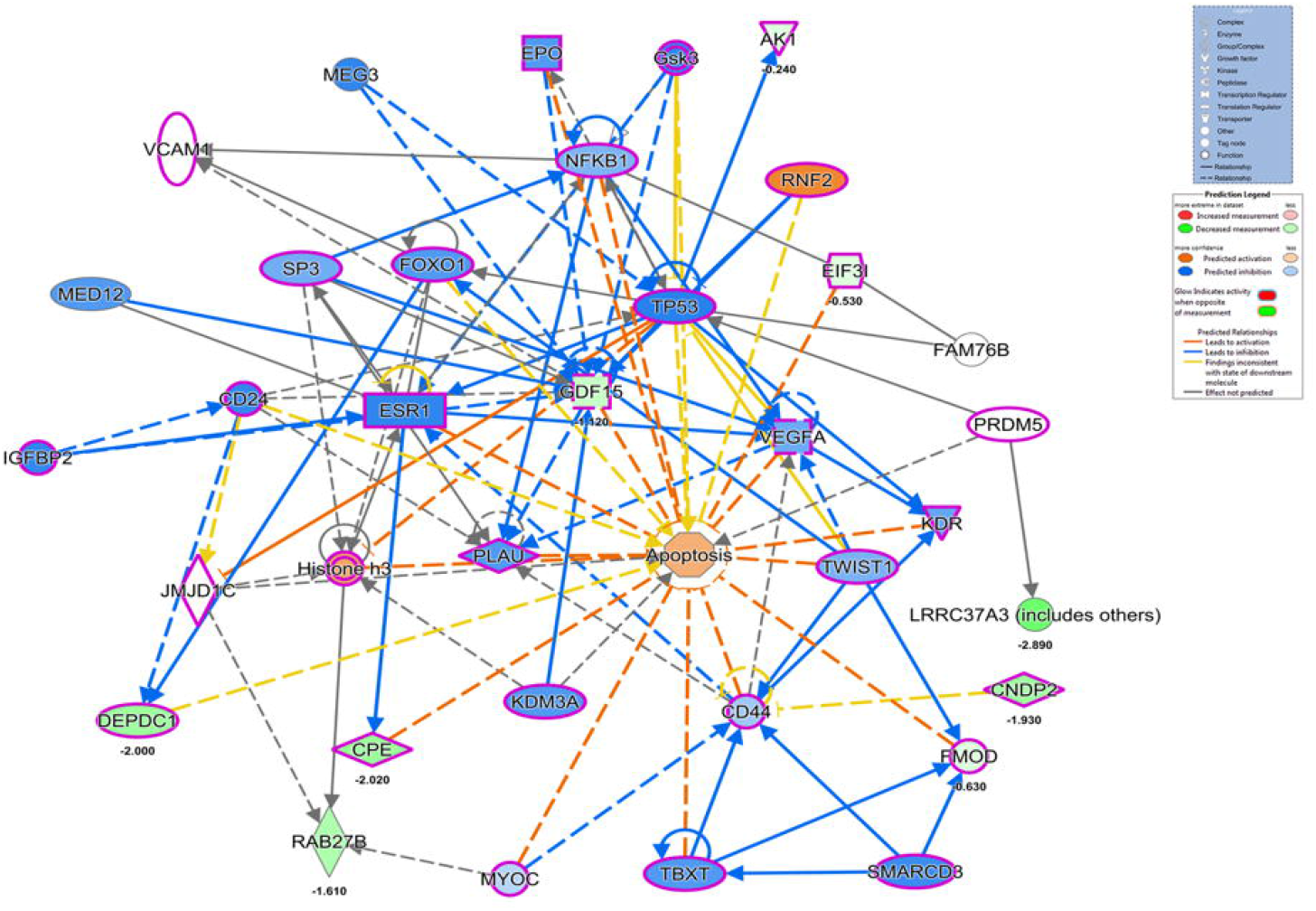
Ingenuity pathway Analysis of underexpressed proteins in Idiopathic infertile patients compared to fertile donor top Disease and functions are Cancer, Cell Death and Survival, Organismal Injury and abnormalities.

IPA canonical pathway revealed that Aryl Hydrocarbon Receptor Signaling, Hypoxia Signaling in the Cardiovascular System, Telomerase Signaling, PPAR Signaling, Xenobiotic Metabolism Signaling, eNOS Signaling and Oxidative Phosphorylation were deregulated **(Fig. 8, Table S6)**

**Fig. 8.**
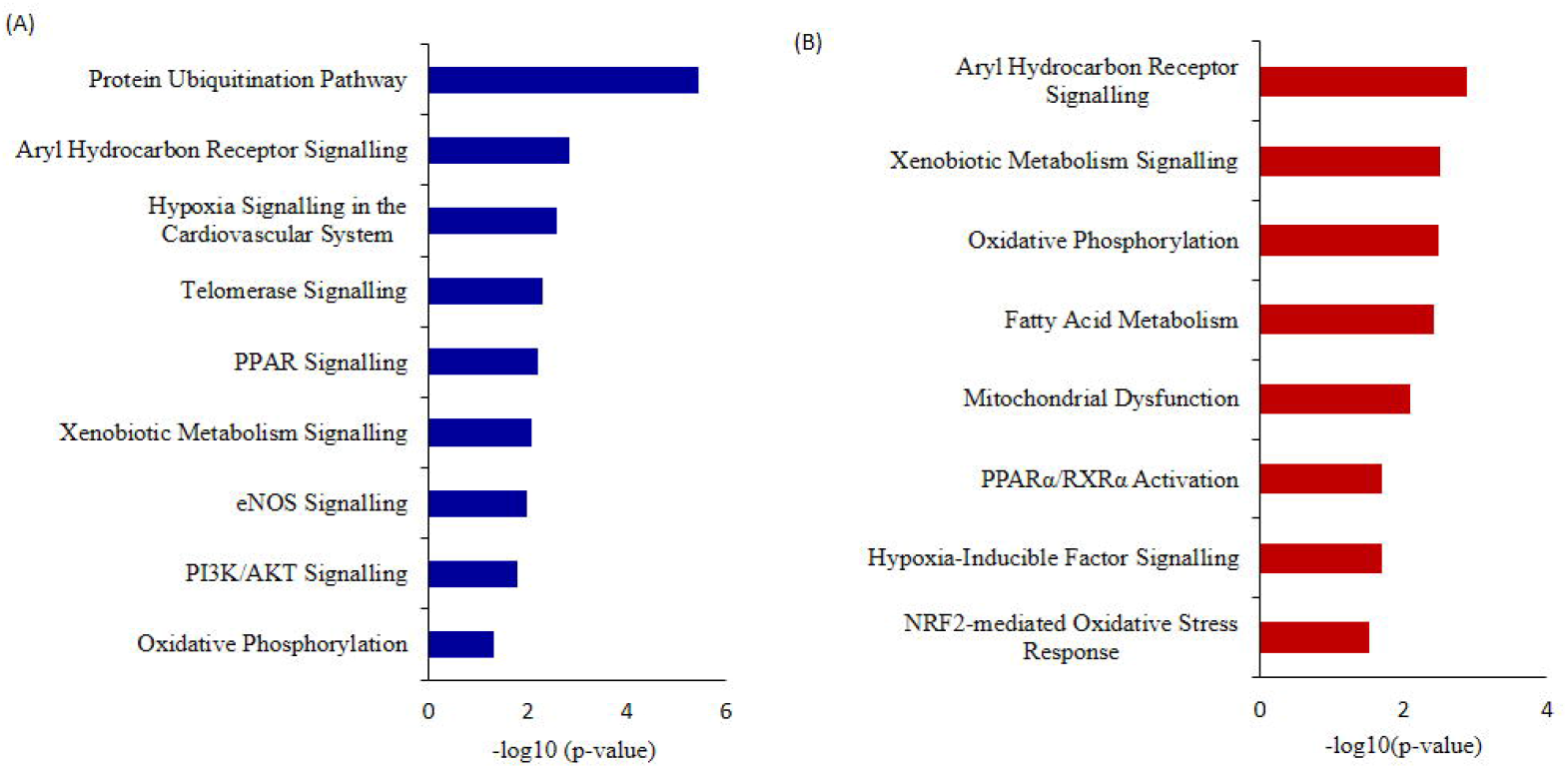
Ingenuity Pathway Analysis (IPA) (A) Canonical pathways analysis and (B) toxicity lists analysis of the differentially expressed proteins (DEPs) of idiopathic infertile patients in comparison to fertile donors.

The top toxicity list and functions determined by IPA-Toxicological pathway showed that Aryl Hydrocarbon Receptor (AhR) Signalling, Xenobiotic Metabolism Signalling, Fatty Acid Metabolism, Hypoxia-Inducible Factor Signalling, NRF2-mediated Oxidative Stress Response, Mitochondrial Dysfunction and Oxidative Phosphorylation are the most affected toxicological functions **(Fig. 8, Table S7**).

### 3.6 Expression profile of key pathway proteins

The key pr**o**teins in top canonical pathway are AhR (predicted) and Heat shock protein (HSP)90β validated by western blot (**Fig. 9 C & D**) which corroborated the LC-MS/MS data. Both the proteins were found to be over expressed in the idiopathic infertile patientsas compared to fertile donor. The expression of 4-Hydoxynonenal (HNE) protein adduct and protein nitrosylation were also found to be overexpressed in idiopathic infertile patients (**Fig. 9 A & B**).

**Fig. 9.**
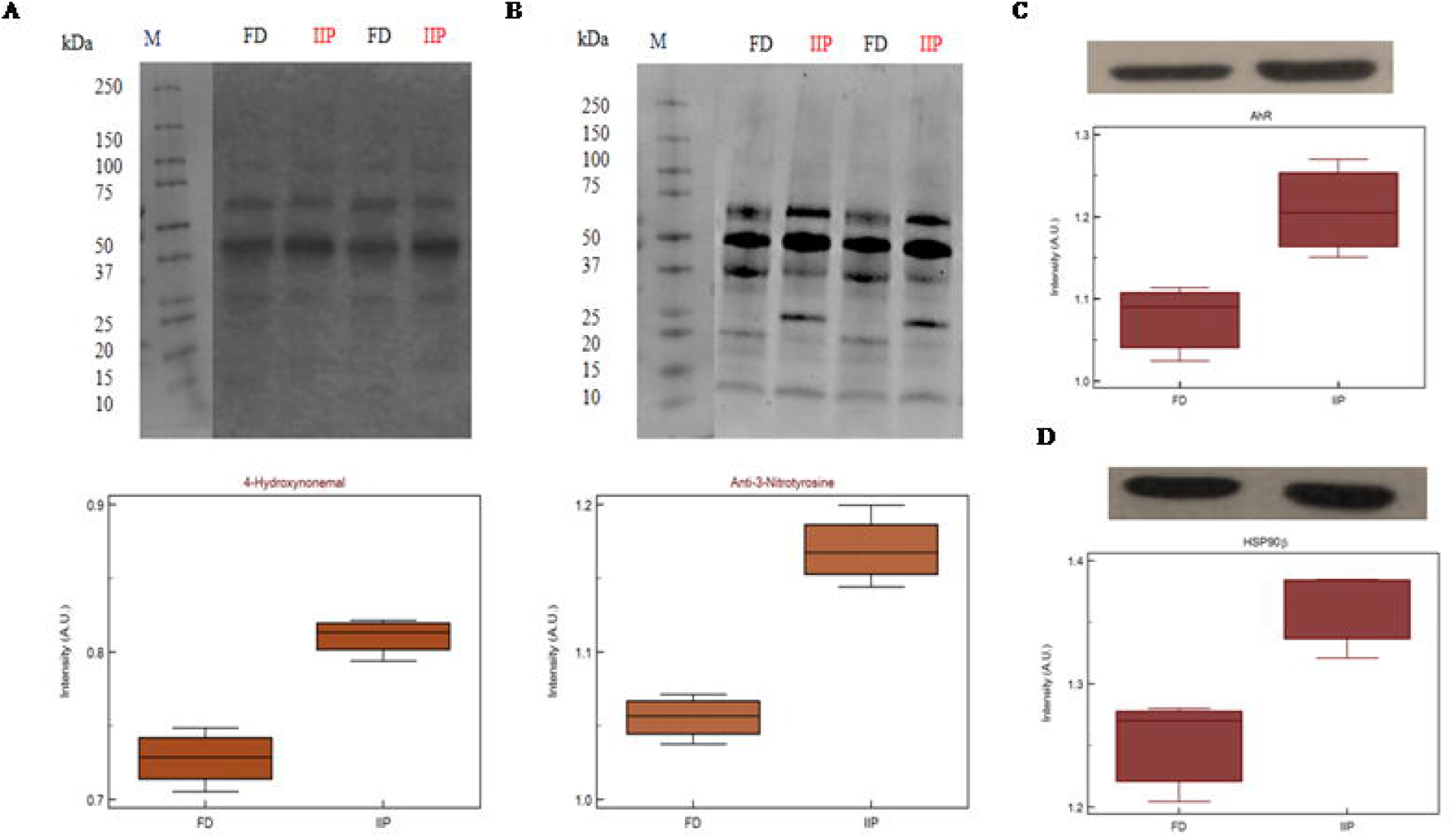
Expression profile of (A) 4-Hydroxynonenal (B) 3-nitrotyrosine (C) AhR (D) HSP90β and their respective densitometry analysis in spermatozoa of FD: fertile donor and IIP: idiopathic infertile patient with total protein normalization (in arbitrary unit). p<0.05 with respect to fertile donor.

## 4. Discussion

Polyaromatic hydrocarbons (PAHs) are known endocrine disruptors which mimic the reproductive hormones and interfere with their synthesis by acting as agonist and antagonists. Several research studies on adult rodents with PAHs such as B(a)P, 2,3,7,8-Tetrachlorodibenzodioxin and 3-Methylchloranthrene results in an increase in the number of abnormal sperm and immature germ cells ^21,22^, affects spermatogenesis by causing testicular atrophy ^23^, diminishes testicular weight and increase apoptosis in seminiferous tubules ^24,25^. Apart from this PAHs also disrupt the normal embryonic development by inducing oxidative stress. A study by Delfino, demonstrated that exposure to PAHs disrupt the redox balance and generate reactive oxygen species (ROS) ^26^. This leads to oxidative stress causing damage to biomolecules such as DNA, lipid and protein involved in the development of reproductive process. In the current study, seminal PAH concentration was measured in the semen of idiopathic infertile patients followed by proteomics of spermatozoa to understand the mechanism(s) by which PAHs elicit male infertility. Several studies have shown that urinary 1-hydroxypyrene (1-OHP) is a good biological index for the occupational exposure assessment of PAHs ^27^. In a study by Xia et al, men with higher urinary concentrations of 1-hydroxypyrene, 2-hydroxyfluorene and sum of all four PAH metabolites (assessed as tertiles) were more likely to have idiopathic male infertility ^27^. In another study, the same authors ^27^reported that higher urinary 1-hydroxypyrene (assessed as quantiles) levels were more likely to have below reference sperm concentration and total sperm count. Similarly, Song et al found a direct correlation between blood concentrations of PAH with sperm motility and a decrease in pregnancy outcome ^28^. But no concrete data were available on semen concentration of PAH with respect to infertility to justify their involvement in spermatogenesis, sperm maturation and sperm function.

This pioneer study reports association between idiopathic male factor infertility with comprehensive screening data of PAHs in the semen resulting in sperm dysfunction through shotgun proteomic analysis. A total of 13 PAHs out of the 16 standards were identified in our sample implying the ubiquitous incidence in non-occupational peoples (only 4 people in the patient group are smokers). The concentrations of all the 13 PAHs detected were significantly higher (p<0.0001) in idiopathic infertile patients with respect to fertile donors. Based on AUC_ROC_ the PAHs having most significant effect on fertility are of the following order Anthracene<benzo(a)pyrene<benzo[b]fluoranthene<Fluoranthene<benzo(a)anthracene<indol (123CD)pyrene<pyrene<naphthalene<dibenzo(AH)anthracene<fluorene<2bromonaphthalene <chrysene<benzo(GH1)perylene. Irrespective of fertility status all analyzed samples possessed naphthalene, albeit at different concentration showing the highest cut-off value of 868ng/ml. On the other hand, the lowest was noticed for bothchrysene and benzo(a)pyrene at 6 ng/ml and benzo(a)pyrene being the ubiquitous one in idiopathic infertile patients distinctively segregating infertile men from their fertile counter parts. Four out of 43 semen samples analyzed in fertile donor (∼9%) showed measurable benzo(a)pyrene (0.35 ± 1.17ng/ml). On the other hand, substantially high level of benzo(a)pyrene (43.37 ± 38.57ng/ml) was detected in all the 60 semen samples analyzed in idiopathic infertile patients. Thus, benzo(a)pyrene can be used as a marker to distinguish infertile men from fertile one with 66.67% sensitivity and 100% specificity at 95% CI (confidence interval).A large cohort study may further substantiate our findings.Though most of the idiopathic infertile males participated in the present study have normal spermiogram, ∼30% have declined motility while∼60% have above normal anomalous spermatozoa. It will not be out of context to mention that prenatal exposure of benzo(a)pyrene to Gclm knockout mice resulted reduction in testicular weight, testicular sperm head counts, epididymal sperm counts, and epididymal sperm motility when analyzed at 10-weeks of age, relative to wild type littermates ^29^.In another study on Mexican workers in a rubber factory with potential occupational exposure to PAHs, impaired spermatogenesis was reported evinced by increased anomalies in sperm concentration, motility and morphology ^30^.In fact, we have observed increased retention of histone proteins (H1-3, H1-4, H1-7, H2BC19P, H2BC11, H2BC12, H2BS1) in the spermatozoa of idiopathic infertile patients implying improper nuclear remodelling ^31^. Furthermore, the gene ontology and protein-protein interaction analysis data (BiNGO, ClueGO, String) reveal that DNA packaging, chromatin assembly and nucleosome assembly is deregulated in patient group. PAHs are also known to cause potential DNA damage in case of idiopathic infertile males ^32^. Interactive metabolites of PAHs may reach the testicles and epididymis forming sperm DNA adducts ^33^. In addition, the compounds resulting from PAH oxidation have the ability to enter oxidation cycles, which increase the generation of ROS and thus cause sperm DNA damage. To corroborate the findings we observed significantly higher 4-HNE and S-Nitrotyrosine levels in the spermatozoa of infertile patients in comparison to their fertile counterparts. Apart from improper compaction and packaging of sperm DNA, themajor alterations in carbohydrate metabolism and active transport across membrane leads to production of dysfunctional spermatozoa. The predicted alteredNADH regeneration pathway further corroborates the imbalance in cellular redox state which is expected from redox acting toxicants like PAHs ^34^.

In this study, 71 DEPs were reported in patient group with higher levels of PAH and IPA toxicity list of these DEPs was predicted to be involved in Aryl Hydrocarbon Receptor (AhR) Signalling, Xenobiotic Metabolism Signalling, Hypoxia-Inducible Factor Signalling, NRF2-mediated Oxidative Stress Response, Mitochondrial Dysfunction and Oxidative Phosphorylation.The AhR is ligand-activated transcription factor that responds to endogenous ligands in addition to exogenous xenobiotic ligands, such as PAHs ^35^. Upon ligand binding, AhR translocates to nucleus where it binds to AhR nuclear transporter (ARNT) and activates xenobiotic metabolizing enzymes: cytochrome P450 (CYP) 1A1, 1A2, and 1B1 for catalyzing oxidative biotransformation of xenobiotics ^36,37^. After biotransformation PAHs generate potential reactive intermediates ^38,39^. In fact, Hansen et al., reported the role of AhR signalling in maintenance of Sertoli cell architecture and resultant spermatogenesis in AhR knockout mice where the poorly remodelled spermatozoa are suggested to be more susceptible to oxidant attack ^40^. In the present study an increased expression of 4-HNE and 3-Nitrotyrosine implies induction of oxidative stress. On the other hand, 4-HNE is known to produce DNA adduct ^41^ as observed in case of PAH exposure and AhR signalling^42^.It is pertinent to mention here that the levels of 4-HNE within spermatozoa are positively correlated with mitochondrial superoxide formation ^43^, and elevated 4-HNE is responsible for numerous adverse effects on sperm function such as decline in motility, morphology, acrosome reaction, sperm-oocyte interaction and apoptosis ^44,45^. The BiNGO and IPA canonical pathway analysis further supports the hypothesis. Protein S-nitrotyrosination is responsible for protection of the proteins under oxidative stress, however irreversible S-nitrotyrosination leads to pathological condition. Of late, it has been elucidated that hydrophobic bio-structures like cell membranes and lipoproteins undergo S-nitrosylation and has strong association with lipid peroxidation^46^. Therefore, it is quite natural to observe an increase in both 4-HNE and 3-nitrotyrosine concentrations in the spermatozoa of infertile patients implying PAH-induced oxidative stress. Further, experimental strategies may reveal the proximal oxidizing mechanism during tyrosine nitration including mapping and identification of the tyrosine nitration sites in specific proteins in the spermatozoa of idiopathic infertile men. Moreover, parallel over-expression of AhR and HSP90β as observed by western blot in the present study corroborated the finding as it is suggested that ligand-bound AhR translocates to the nucleus with HSP90β showing its co-localization in the nucleus ^47^. In contrast to AhR-dependent and CYP1A-mediated production of intracellular ROS, the AhR signaling pathway also regulates the expression of genes involved in antioxidant responses. Besides AhR signaling, NRF2 is another important transcription factor regulating genes that are critically involved in the metabolism of xenobiotics as well as endogenous compounds which is reported as top toxicological pathway in our DEP data set. Both signaling pathways respond to environmental and endogenous stressors. Albeit, AhR and NRF2 are clearly separated signaling pathways, recent reports demonstrate the cross-regulation between these two signaling axes suggesting an integrated response to environmental stressors ^48^.

Glutathione, an important antioxidant is involved in the elimination of PAHs and spermatozoa depend heavily on glutathione metabolizing system for its survival^49^. The reactive intermediates formed after metabolism of PAHs are conjugated to glutathione and eliminated by glutathione-S-transferases (GST) and glutathione peroxidase (GPx) ^50^. Glutathione (GSH) acts as a redox sensor by oxidizing to glutathione disulfide molecule (GSSG). So the ratio of GSH to GSSG is used as a biosignature of oxidized intracellular environment ^51^.A recent report by Branco et al., 2021 have reported that PAH and their metabolites show idiosyncratic behaviour with respect to glutathione metabolism where phenanthrene induced higher ROS production. On the other hand, the authors reported increased GSH levels by benzo(b)fluoranthene along with augmented levels of protein sulfydryl group. The upregulation of GSH was opined to be a consequence of Nrf2 signalling activation and increased levels of glutathione metabolising enzymes and their mRNA after exposure to benzo(b)fluoranthene, but not during exposure to phenanthrene ^52^.Moreover, data of the present study shows a distinct energy deprived and hypoxic state in the spermatozoa of infertile men due to declined redox potential which is similar to the results of previously report by our group for unilateral varicocele patients ^53^.

## Conclusion

The present findings surmise the adverse impact of environment borne PAHs exposure on sperm function in idiopathic infertility which are largely ignored in regular infertility assessment. The high level of benzo(a)pyrene in the infertile group could serve as a predictive marker for idiopathic infertility along with the signature proteins AhR and the HSPs, particularly the HSP90. The presence of oxidative protein modification and differential expression of proteins involved in chromatin packaging and DNA damage further corroborates the noxious effect of PAH in semen (Figure 10). Therefore, it is suggested that along with seminogram and other biological markers, analysis of seminal levels of environmental toxins such as PAH in general and benzo(a)pyrene in particular may help in proper management of idiopathic male factor infertility.

**Fig. 10.**
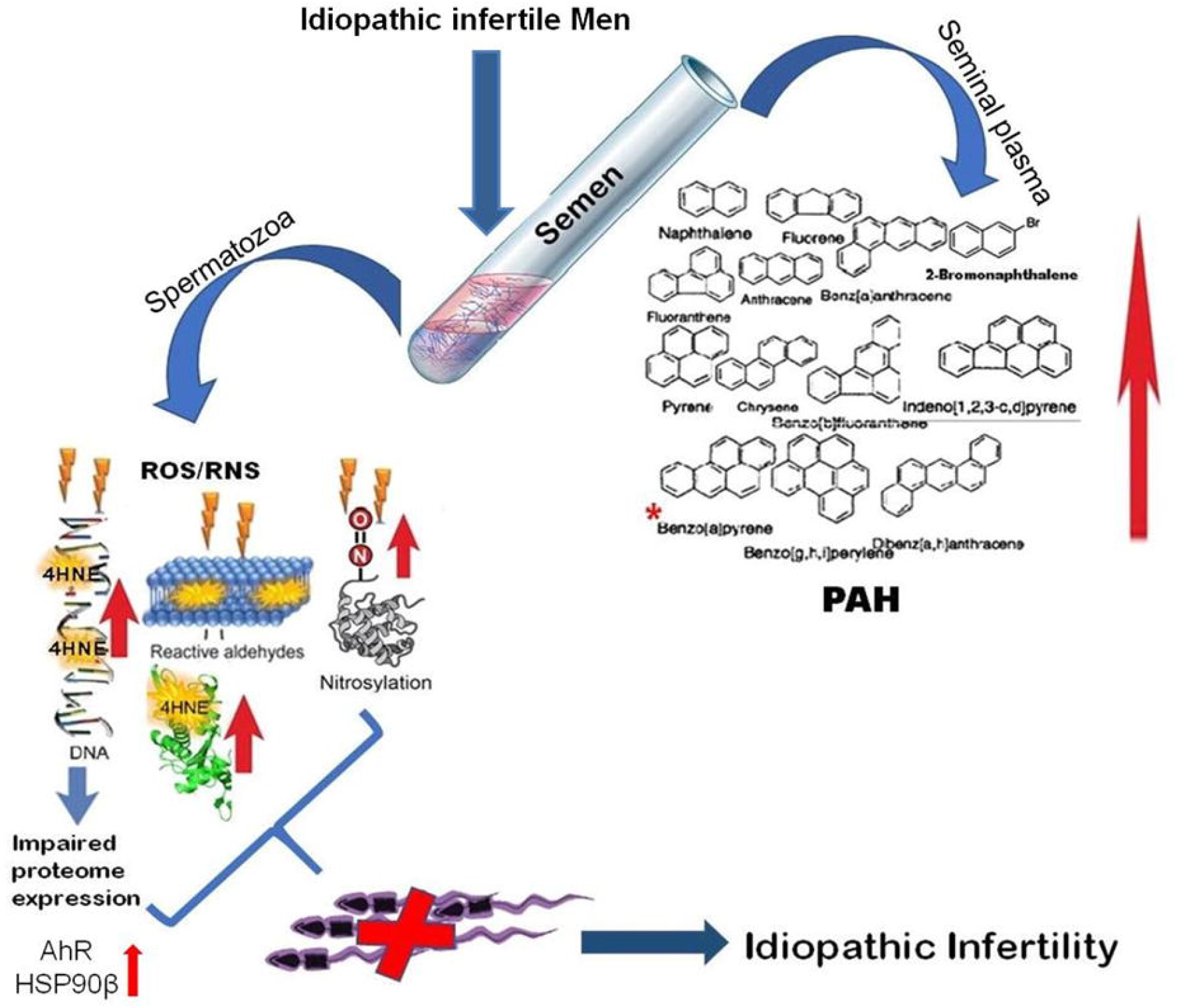
Schematic representation of molecular mechanisms involved in environmental borne polyaromatic hydrocarbon induced sperm dysfunction in idiopathic male infertility.

## Acknowledgement

The authors thank Director, DBT-Institute of Life Science, Bhubaneswar, India for the computational facilities.

