## Supplemental Table 1 for "Association of seminal polyaromatic hydrocarbons exposome with idiopathic male factor infertility: A proteomic insight into sperm function"

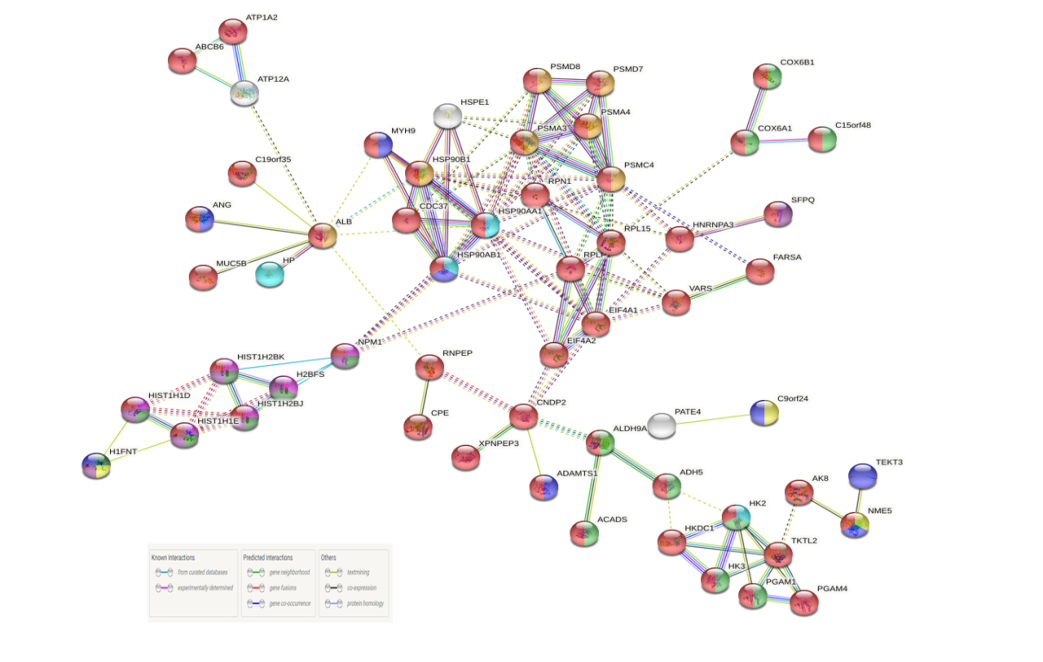

**Fig. 1:** String protein-protein interaction of differentially expressed proteins (DEPs) with MCL clustering having a confidence threshold 0.04.

**Table S1:** Semen parameters in fertile donor and Idiopathic infertile patients.

| **Parameter** | **WHO** | **Fertile donor** | **Idiopathic infertile patient** | **p-value** |
| --- | --- | --- | --- | --- |
| Volume (mL) | >1.5 | 3.24± 0.49 | 3.03± 0.64 | P = 0.0750 |
| pH | 7.2-8.0 | 7.39 ± 0.18 | 7.35± 0.17 | P = 0.2806 |
| Sperm Motility (%) | >40 | 56.30± 8.40 | 43.42± 11.37 | P < 0.0001 |
| Sperm Concentration (10^6^/mL) | >15 | 123.33± 22.30 | 86.77± 40.88 | P < 0.0001 |
| Sperm viability (%) | > 58 | 53.45 ± 10.23 | 39.75 ± 9.45 | P < 0.0001 |
| Sperm morphology (%) | > 4 | 12.5 ± 1.5 | 20.5 ± 5.3 | P < 0.0001 |

**Table S2:** Receiver Operating Characteristic (ROC) optimal cut points and operating characteristics for semen PAH metabolites classification of fertile versus Idiopathic infertile patients.

| **Sl. No.** | **PAH** | **Area Under the ROC Curve** | **Standard Error** | **95% Confidence Interval** | **Sensitivity** | **Specificity** | **Youden Index J** | **Cut-off Value** | **p Value** |
| --- | --- | --- | --- | --- | --- | --- | --- | --- | --- |
| 1 | 2-Bromonapthalene | 0.688760 | 0.0419 | 0.589983 to 0.776351 | 50.00 | 88.37 | 0.3837 | >188 | <0.0001 |
| 2 | Benzo(a)Pyrene | 0.817829 | 0.0344 | 0.729595 to 0.887001 | 66.67 | 100.00 | 0.6667 | >6 | <0.0001 |
| 3 | Anthracene | 0.822093 | 0.0365 | 0.734391 to 0.890464 | 70.00 | 100.00 | 0.7000 | >62 | <0.0001 |
| 4 | Benzo(a)Anthracene | 0.742829 | 0.0388 | 0.647304 to 0.823913 | 53.33 | 100.00 | 0.5333 | >333 | <0.0001 |
| 5 | Benzo(b)Fluoranthene | 0.793798 | 0.0362 | 0.702822 to 0.867211 | 63.33 | 97.67 | 0.6101 | >40 | <0.0001 |
| 6 | Benzo(GH1)Perylene | 0.652326 | 0.0356 | 0.552144 to 0.743496 | 35.00 | 100.00 | 0.3500 | >107 | <0.0001 |
| 7 | Chrysene | 0.686047 | 0.0348 | 0.587144 to 0.773925 | 40.00 | 100.00 | 0.4000 | >6 | <0.0001 |
| 8 | Dibenzo(AH)Anthracene | 0.710078 | 0.0337 | 0.612407 to 0.795283 | 43.33 | 100.00 | 0.4333 | >7 | <0.0001 |
| 9 | Fluoranthene | 0.744186 | 0.0370 | 0.648762 to 0.825086 | 51.67 | 100.00 | 0.5167 | >17 | <0.0001 |
| 10 | Fluorene | 0.690310 | 0.0375 | 0.591607 to 0.777735 | 43.33 | 100.00 | 0.4333 | >831 | <0.0001 |
| 11 | Indo(123CD)Pyrene | 0.725969 | 0.0384 | 0.629270 to 0.809246 | 45.00 | 100.00 | 0.4500 | >8 | <0.0001 |
| 12 | Napthalene | 0.718992 | 0.0459 | 0.621851 to 0.803132 | 58.33 | 100.00 | 0.5833 | >868 | <0.0001 |
| 13 | Pyrene | 0.732558 | 0.0407 | 0.636300 to 0.814997 | 55.00 | 97.67 | 0.5267 | >32 | <0.0001 |

**Table S3:** List of differentially expressed proteins in idiopathic infertile patients as compared to fertile donor.

| **Sl No** | **ID** | **Gene name** | **Symbol** | **Log_2_FC** | **p-value** |
| --- | --- | --- | --- | --- | --- |
| 1 | P14854 | Cytochrome c oxidase subunit 6B1 | COX6B1 | -4.46 | 0.028 |
| 2 | P0C8F1 | Prostate and testis expressed 4 | PATE4 | -2.99 | 0.028 |
| 3 | A6NM11 | Leucine rich repeat containing 37 member A3 | LRRC37A3 | -2.89 | 0.044 |
| 4 | P04843 | Ribophorin I | RPN1 | -2.42 | 0.045 |
| 5 | P16870 | Carboxypeptidase E | CPE | -2.02 | 0.011 |
| 6 | Q5TB30 | DEP domain containing 1 | DEPDC1 | -2 | 0.008 |
| 7 | Q96KP4 | Carnosinedipeptidase 2 | CNDP2 | -1.93 | 0.014 |
| 8 | Q9UHI8 | Apolipoprotein E | APOE | -1.7 | 0.042 |
| 9 | O00194 | RAB27B, member RAS oncogene family | RAB27B | -1.61 | 0.015 |
| 10 | P61604 | Heat shock protein family E (Hsp10) member 1 | HSPE1 | -1.5 | 0.001 |
| 11 | Q8N0Y7 | Phosphoglycerate mutase family member 4 | PGAM4 | -1.43 | 0.021 |
| 12 | Q99988 | Growth differentiation factor 15 | GDF15 | -1.12 | 0.039 |
| 13 | P12074 | Cytochrome c oxidase subunit 6A1 | COX6A1 | -1.04 | 0.026 |
| 14 | O14618 | Copper chaperone for superoxide dismutase | CCS | 1.01 | 0.031 |
| 15 | P56597 | NME/NM23 family member 5 | NME5 | 1.03 | 0.038 |
| 16 | P03950 | Angiogenin | ANG | 1.15 | 0.040 |
| 17 | P51665 | Proteasome 26S subunit, non-atpase 7 | PSMD7 | 1.18 | 0.003 |
| 18 | P08238 | heat shock protein 90 alpha family class B member 1 | HSP90AB1 | 1.2 | 0.018 |
| 19 | Q6ZQR2 | Cilia and flagella associated protein 77 | CFAP77 | 1.2 | 0.017 |
| 20 | P14625 | Heat shock protein 90 beta family member 1 | HSP90B1 | 1.2 | 0.045 |
| 21 | Q8IYK2 | Coiled-coil domain containing 105 | CCDC105 | 1.21 | 0.011 |
| 22 | Q5JNZ5 | Ribosomal protein S26 pseudogene 11 | RPS26P11 | 1.21 | 0.018 |
| 23 | P07900 | Heat shock protein 90 alpha family class A member 1 | HSP90AA1 | 1.3 | 0.039 |
| 24 | Q8IZ16 | Chromosome 7 open reading frame 61 | C7orf61 | 1.41 | 0.007 |
| 25 | P18669 | Phosphoglycerate mutase 1 | PGAM1 | 1.43 | 0.031 |
| 26 | Q9NQH7 | X-prolyl aminopeptidase 3 | XPNPEP3 | 1.43 | 0.016 |
| 27 | Q9H4A4 | Arginyl aminopeptidase | RNPEP | 1.45 | 0.001 |
| 28 | Q08648 | Sperm associated antigen 11B | SPAG11B | 1.46 | 0.004 |
| 29 | P11766 | Alcohol dehydrogenase 5 (class III), chi polypeptide | ADH5 | 1.48 | 0.005 |
| 30 | Q96MA6 | Adenylate kinase 8 | AK8 | 1.49 | 0.020 |
| 31 | Q99417 | MYC binding protein | MYCBP | 1.5 | 0.034 |
| 32 | Q75WM6 | H1.7 linker histone | H1-7 | 1.58 | 0.016 |
| 33 | Q6DRA6 | H2B clustered histone 19, pseudogene | H2BC19P | 1.58 | 0.022 |
| 34 | Q9Y285 | Phenylalanyl-trnasynthetase subunit alpha | FARSA | 1.6 | 0.012 |
| 35 | P16219 | Acyl-Coa dehydrogenase short chain | ACADS | 1.72 | 0.015 |
| 36 | P26640 | Valyl-trnasynthetase 1 | VARS | 1.72 | 0.033 |
| 37 | Q9HC84 | Mucin 5B, oligomeric mucus/gel-forming | MUC5B | 1.76 | 0.006 |
| 38 | P52789 | Hexokinase 2 | HK2 | 1.82 | 0.008 |
| 39 | P52790 | Hexokinase 3 | HK3 | 1.82 | 0.034 |
| 40 | Q2TB90 | Hexokinase domain containing 1 | HKDC1 | 1.82 | 0.034 |
| 41 | Q6ZS72 | PEAK family member 3 | PEAK3 | 1.82 | 0.047 |
| 42 | P23246 | Splicing factor proline and glutamine rich | SFPQ | 1.82 | 0.033 |
| 43 | Q969V4 | Tektin 1 | TEKT1 | 1.85 | 0.006 |
| 44 | Q9BXF9 | Tektin 3 | TEKT3 | 1.87 | 0.041 |
| 45 | Q14990 | Outer dense fiber of sperm tails 1 | ODF1 | 2 | 0.028 |
| 46 | P54707 | Atpase H+/K+ transporting non-gastric alpha2 subunit | ATP12A | 2.04 | 0.043 |
| 47 | P50993 | Atpase Na+/K+ transporting subunit alpha 2 | ATP1A2 | 2.04 | 0.019 |
| 48 | P60842 | Eukaryotic translation initiation factor 4A1 | EIF4A1 | 2.05 | 0.008 |
| 49 | Q14240 | Eukaryotic translation initiation factor 4A2 | EIF4A2 | 2.05 | 0.008 |
| 50 | Q9BR76 | Coronin 1B | CORO1B | 2.08 | 0.011 |
| 51 | P02768 | Albumin | ALB | 2.09 | 0.030 |
| 52 | P06899 | H2B clustered histone 11 | H2BC11 | 2.1 | 0.017 |
| 53 | O60814 | H2B clustered histone 12 | H2BC12 | 2.1 | 0.017 |
| 54 | P57053 | H2B.S histone 1 | H2BS1 | 2.1 | 0.017 |
| 55 | P35579 | Myosin heavy chain 9 | MYH9 | 2.1 | 0.036 |
| 56 | P00738 | Haptoglobin | HP | 2.16 | 0.022 |
| 57 | Q6ZNM6 | Testis expressed 43 | TEX43 | 2.2 | 0.032 |
| 58 | Q9C002 | Chromosome 15 open reading frame 48 | C15orf48 | 2.24 | 0.014 |
| 59 | Q14093 | Cylicin 2 | CYLC2 | 2.32 | 0.012 |
| 60 | P51991 | Heterogeneous nuclear ribonucleoprotein A3 | HNRNPA3 | 2.41 | 0.032 |
| 61 | P61313 | Ribosomal protein L15 | RPL15 | 2.44 | 0.035 |
| 62 | A6NJV1 | Family with sequence similarity 166 member C | FAM166C | 2.52 | 0.035 |
| 63 | Q8NCR6 | Chromosome 9 open reading frame 24 | C9orf24 | 2.63 | 0.038 |
| 64 | P06748 | Nucleophosmin 1 | NPM1 | 2.9 | 0.026 |
| 65 | O75610 | Left-right determination factor 1 | LEFTY1 | 2.94 | 0.045 |
| 66 | Q9NP58 | ATP binding cassette subfamily B member 6 (Langereis blood group) | ABCB6 | 3 | 0.007 |
| 67 | P49189 | Aldehyde dehydrogenase 9 family member A1 | ALDH9A1 | 3.04 | 0.000 |
| 68 | P16402 | H1.3 linkher histone, cluster member | H1-3 | 3.32 | 0.010 |
| 69 | P10412 | H1.4 linker histone, cluster member | H1-4 | 3.32 | 0.009 |
| 70 | Q9H0I9 | Transketolase like 2 | TKTL2 | 3.42 | 0.004 |
| 71 | P05386 | Ribosomal protein lateral stalk subunit P1 | RPLP1 | 4.12 | 0.004 |

**Table S4:** Enriched gene ontology of differentially expressed proteins using ClueGO app with their cellular localization, molecular functions and biological process inidiopathic infertile patients as compared to fertile donor.

| **GOID** | **GOTerm** | **log 10p-value** | **% Associated Genes** | **Associated Genes Found** |
| --- | --- | --- | --- | --- |
| **Cellular component** | |  |  |  |
| GO:0098533 | ATPase dependent transmembrane transport complex | 4.113085 | 12.5 | [ABCB6, ATP12A, ATP1A2] |
| GO:0036126 | sperm flagellum | 4.147414 | 4.032258 | [AK8, NME5, ODF1, PGAM4, TEKT3] |
| GO:0071682 | endocytic vesicle lumen | 4.169997 | 13.04348 | [HP, HSP90AA1, HSP90B1] |
| GO:0000786 | nucleosome | 4.395893 | 4.545455 | [H1-3, H1-4, H2BC11, H2BC12, H2BC19P] |
| **Molecular functions** | |  |  |  |
| GO:0008235 | metalloexopeptidase activity | 3.983048 | 5.714286 | [CNDP2, CPE, RNPEP, XPNPEP3] |
| GO:0004396 | hexokinase activity | 4.002299 | 11.53846 | [HK2, HK3, HKDC1] |
| GO:0005536 | glucose binding | 4.353875 | 15 | [HK2, HK3, HKDC1] |
| **Biological process** | |  |  |  |
| GO:0030261 | chromosome condensation | 3.215314 | 6 | [H1-3, H1-4, H1-7] |
| GO:0004396 | hexokinase activity | 4.068022 | 11.53846 | [HK2, HK3, HKDC1] |
| GO:0006334 | nucleosome assembly | 5.024906 | 4.054054 | [H1-3, H1-4, H2BC11, H2BC12, H2BC19P, NPM1] |
| GO:0061621 | canonical glycolysis | 5.08326 | 10.25641 | [HK2, HK3, PGAM1, PGAM4] |
| GO:0006735 | NADH regeneration | 5.08326 | 10.25641 | [HK2, HK3, PGAM1, PGAM4] |

**Table S5:** Enriched disease and function network of differentially expressed proteins in Idiopathic Infertile patients compared to fertile donor.

| **Sl. No.** | **Molecules in Network** | **Score** | **Focus Molecules** | **Top Diseases and Functions** |
| --- | --- | --- | --- | --- |
| **A** | **Upregulated DEPs** |  |  |  |
| 1 | ABCB6,ADH5,AHR,AIP,ANG,ATP1A2,BMI1,CDK4,EGLN,ERK1/2,GRB7,GRK5,H14,HK2,HKDC1,Hsp90,HSP90AA1,HSP90AB1,**HSP90B1**,LIMCH1,mir-128,MYCBP,MYH9,myosin-light-chain kinase,NPM1,PGAM1,PI3K p85,PRINS,PSMD7,RNASE4,RPL13A,SBDS,SFPQ,SMARCA4,TRAP1 | 35 | 18 | [Cancer, Endocrine System Disorders, Organismal Injury and Abnormalities] |
| 2 | AGT,ALB,APCS,ATF1,Cbp/p300,CCS,CD163,CREB1,CRYAB,CYBA,EIF4A1,FST,GPS2,GRB2,H1-3,H19,H2BC12,HDAC3,HIF1A,HNRNPA3,HP,IL22,LEFTY1,Mek,MEN1,MKNK1,MUC1,MUC5B,Nr1h,RNASE1,RPL15,TBL1XR1,TCF4,TCF7L2,TNF | 21 | 12 | [Cell Death and Survival, Cellular Development, Organismal Survival] |
| **B** | **Downregulated DEPs** |  |  |  |
| 3 | AK1,CD24,CD44,CNDP2,CPE,DEPDC1,EIF3I,EPO,ESR1,FAM76B,FMOD,FOXO1,GDF15,Gsk3,Histone h3,IGFBP2,JMJD1C,KDM3A,KDR,LRRC37A3 (includes others),MED12,MEG3,MYOC,NFKB1,PLAU,PRDM5,RAB27B,RNF2,SMARCD3,SP3,TBXT,TP53,TWIST1,VCAM1,VEGFA | 19 | 9 | [Cancer, Cell Death and Survival, Organismal Injury and Abnormalities] |

**Table S6:**Ingenuity Canonical Pathways of the differentially expressed proteins inidiopathic infertile patients as compared to fertile donor along with their significance values and associated molecules.

| **Sl No** | **Ingenuity Canonical Pathways** | **-log (p-value)** | **Ratio** | **Molecules** |
| --- | --- | --- | --- | --- |
| 1 | Protein Ubiquitination Pathway | 5.45 | 0.0444 | CRYAB, HSP90AA1, HSP90AB1, HSP90B1, HSPE1, ODF1, PSMA6, PSMD7 |
| 2 | Aryl Hydrocarbon Receptor Signalling | 2.83 | 0.0421 | ALDH9A1, HSP90AA1, HSP90AB1, HSP90B1 |
| 3 | Hypoxia Signalling in the Cardiovascular System | 2.6 | 0.0577 | HSP90AA1, HSP90AB1, HSP90B1 |
| 4 | Telomerase Signalling | 2.31 | 0.0455 | HSP90AA1, HSP90AB1, HSP90B1 |
| 5 | PPAR Signalling | 2.2 | 0.0417 | HSP90AA1, HSP90AB1, HSP90B1 |
| 6 | Xenobiotic Metabolism Signalling | 2.07 | 0.026 | ALDH9A1, HSP90AA1, HSP90AB1, HSP90B1 |
| 7 | eNOS Signalling | 2 | 0.0353 | HSP90AA1, HSP90AB1, HSP90B1 |
| 8 | Oxidative Phosphorylation | 1.32 | 0.0299 | COX6A1,COX6B1 |
| 9 | PI3K/AKT Signalling | 1.79 | 0.0294 | HSP90AA1, HSP90AB1, HSP90B1 |

**Table S7 :** Ingenuity Toxicity Lists predicted by IPA-Tox for a toxicity analysis along with their significance values and associated molecules

| **Sl. No.** | **Ingenuity Toxicity Lists** | **-log (p-value)** | **Ratio** | **Molecules** |
| --- | --- | --- | --- | --- |
| 1 1 | Aryl Hydrocarbon Receptor Signalling | 2.88 | 0.0268 | ALDH9A1, HSP90B1, HSP90AB1, HSP90AA1 |
| 2 | Xenobiotic Metabolism Signalling | 2.51 | 0.0162 | ALDH9A1, HSP90B1, ADH5, HSP90AB1, HSP90AA1 |
| 3 | Fatty Acid Metabolism | 2.42 | 0.0306 | ALDH9A1, ACADS, ADH5 |
| 4 | PPARα/RXRα Activation | 1.7 | 0.0167 | HSP90B1, HSP90AB1, HSP90AA1 |
| 5 | Hypoxia-Inducible Factor Signalling | 1.69 | 0.029 | HSP90AB1, HSP90AA1 |
| 6 | NRF2-mediated Oxidative Stress Response | 1.52 | 0.0142 | HSP90B1, HSP90AB1, HSP90AA1 |
| 7 | Mitochondrial Dysfunction | 2.09 | 0.0121 | COX6A1,COX6B1 |
| 8 | Oxidative Phosphorylation | 2.48 | 0.0192 | COX6A1,COX6B1 |
